# Neonatal brain-age models in full- and preterm infants

**DOI:** 10.64898/2026.01.23.701337

**Authors:** Howard Chiu, Adam C. Richie-Halford, Molly F. Lazarus, Ariel Rokem, Rocio Velasco Poblaciones, Virginia A. Marchman, Katherine E. Travis, Melissa L. Scala, Heidi M. Feldman, Jason D. Yeatman

## Abstract

Prematurity affects brain development and increases risk for neurodevelopmental impairments. Yet reliable biomarkers for at-risk infants remain limited. The goals of this study are (1a) to construct a white matter neonatal brain-age model including full-term and preterm neonates from a large publicly-available data set, (1b) to evaluate the accuracy of this model for characterizing the preterm brain from the same data set; (2) to determine if a similar model can predict brain-age based on clinical MRI scans from high-risk neonates born preterm; and (3) to evaluate whether this predictive model provides information about the infant’s health beyond conventional clinical and demographic measures. We developed brain-age prediction models using diffusion magnetic resonance imaging-derived white matter features from two datasets: (1) the developing Human Connectome Project (dHCP; 368 healthy infants) and (2) a clinical sample collected at the Lucile Packard Children’s Hospital (LPCH; 162 high-risk preterm infants). White matter features demonstrated strong predictive performance in the dHCP dataset (within 3.9 days) and the LPCH clinical dataset (within 6.6 days). However, brain-age metrics (i.e., brain-age gap) showed no significant associations with health complications measured by a composite score of common prematurity complications. While tractometry-derived brain-age models accurately characterize brain maturation in the neonatal brain, their sensitivity to clinical complications in preterm infants appears limited. Global white matter maturation measures derived from clinical grade data may be insufficiently sensitive to capture the cumulative burden of prematurity-related morbidities, suggesting need for multimodal or longitudinal biomarkers.

**HIGHLIGHTS:** Brain-age models of neonatal dMRI data predict age within days.

Brain-age models of preterm infants predict post-menstrual age within a week.

Brain-age models are accurate with research and clinical dMRI data in preterm infants.

Brain-age gap was not sensitive to health status in a clinical preterm sample.

**GRAPHICAL ABSTRACT:** 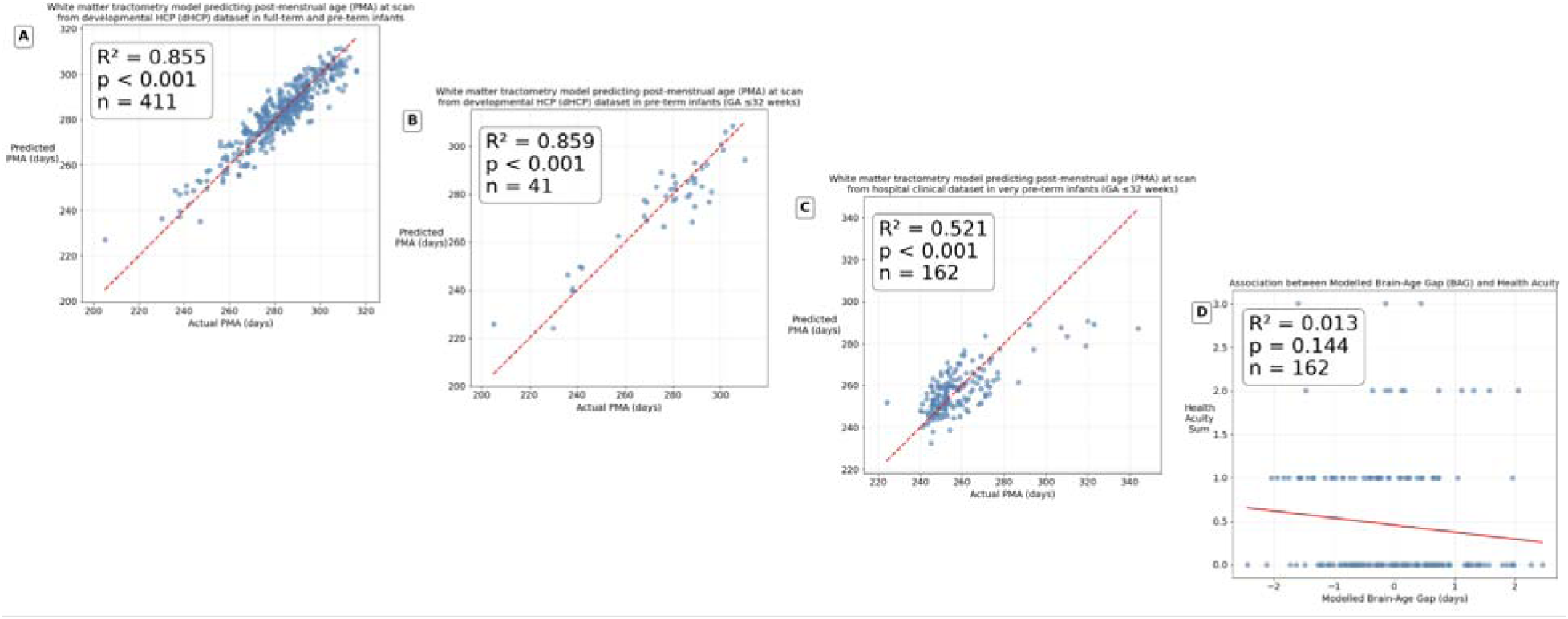

## INTRODUCTION

Prematurity, defined as birth before 37 weeks of gestation, presents a significant global health challenge. According to the World Health Organization, in 2020, the global rate of preterm birth ranged from 4% to 16% (Spoto et al., 2024). Improvements in postnatal care in delivery rooms and neonatal intensive care units have resulted in improvements in mortality rates without changes in morbidity rates (Behrman & Stith Butler, 2007). Thus, while more infants survive, the proportion of survivors with poor neurodevelopmental and cognitive outcomes has largely remained unchanged.

Children born preterm, especially those born before 32 weeks gestation (very preterm; VPT), show heightened risk of long-term neurodevelopmental impairments affecting motor, cognitive, language, and social-emotional functions. Children born VPT have higher rates of grade repetition and increased need of special education services compared to children born at term (Aarnoudse-Moens et al., 2009; Bhutta et al., 2002; McBryde et al., 2020; Moore et al., 2012). On average, preterm-born children score half a standard deviation lower than their term-born peers in standardized assessments of math and reading (McBryde et al., 2020).

An important factor in the pathogenesis of adverse outcomes is the immature brain of the VPT infant, which is particularly vulnerable to various insults, including hypoxia, ischemia, and inflammation. White matter injury, particularly diffuse injuries that are distributed throughout the white matter, is the most prevalent consequence of these insults, owing to the vulnerability of early differentiating preoligodendrocytes, the cell line that in maturity creates myelin. The period from 23 to 32 weeks’ gestation constitutes the period of highest risk for white matter injury, peaking at 28 weeks’ gestation (Inder et al., 2023). Preterm birth can disrupt this critical phase of white matter development, leading to diffuse or focal white matter injury to pre-oligodendrocytes (Inder et al., 2023), which has been associated with neurodevelopmental outcomes (Kline et al., 2021; Roychaudhuri et al., 2024).

Diffusion MRI, which measures the rate and directionality of water diffusion, has shown promise for detecting various white matter injuries that result from prematurity. For example, preterm infants consistently exhibit lower fractional anisotropy (FA) values (an index of the degree of directionality of water diffusion in the white matter) and higher mean diffusivity (MD) values (an index of the average rate of water diffusion) compared to their term counterparts (Dibble et al., 2021; Wu et al., 2020). However, despite the growing evidence that microstructural properties of the premature white matter is relevant to health outcomes, less progress has been made in creating clinically useful models based on diffusion MRI data. One of the challenges has been summarizing white matter development into a single (or a few) metrics that can be used in clinical practice.

The concept of "brain-age" has emerged as a promising biomarker in neuroscience, representing the biological maturity of an individual’s brain as estimated from neuroimaging data (Azzam et al., 2025; Cole et al., 2019; Franke et al., 2010; Smith et al., 2019). Brain-age models are typically constructed using machine learning algorithms trained on neuroimaging data from healthy individuals to predict their chronological age. For children with potential brain injury, the brain-age assigned to an individual child can be compared to the chronological age of the child at the time of scanning. Any difference is labeled a "brain-age gap" or "brain-age index". Brain-age gaps have been established as useful biomarkers in the study of neurological disorders including Parkinson’s (Eickhoff et al., 2021), Alzheimer’s (Franke et al., 2010), traumatic brain injury (Cole et al., 2015), schizophrenia (Constantinides et al., 2023), and major depressive disorder (Han et al., 2021). In pediatric populations, more severe ADHD symptoms in a group of 11-year-olds were significantly associated with younger appearing brains (Kurth et al., 2022), demonstrating the applicability of the brain-age gap across the lifespan.

In the context of neonates, particularly neonates born preterm, brain-age should be trained using postmenstrual age (PMA) at scan as the target variable, rather than chronological age at scan, with the resulting "brain-age gap" derived as the difference between predicted and actual PMA. PMA of a premature infant is defined as the sum of time in utero (gestational age) and time since birth (chronological age), and using PMA ensures that the brain-age gap reflects deviations in the expected trajectory of brain maturation that might be disrupted by premature birth (Ansari et al., 2024; Liu et al., 2023; Sun et al., 2024; Tang et al., 2023). A negative brain-age gap suggests delayed maturation, while a positive gap may indicate accelerated development, and the magnitude of the gap may be related to potential health complications. If brain-age models are sensitive to health status or outcomes they could provide a promising tool for personalizing treatment strategies in neonatal intensive care units by providing information above and beyond clinical data such as birth weight and medical history (Hand et al., 2020; Stevenson et al., 2020).

This retrospective study has three goals: (1a) to construct a neonatal brain-age model including full-term and preterm neonates from a large data set, specifically utilizing high-dimensional white matter features computed via tractometry and then (1b) to evaluate the accuracy of this model for characterizing maturation of the preterm brain on a held-out sample from the same data set; (2) to determine if a neonatal brain-age model could be constructed in the same manner on a new dataset that consists of routine clinical MRI from high-risk neonates born preterm who are scanned on a clinical scanner as standard of practice; and (3) to evaluate whether this preterm white matter-based brain-age model provides additional information about the infant’s health outcomes beyond conventional clinical and demographic measures. We hypothesize that tractometry-derived features will enable accurate brain-age prediction, exceeding the contribution of sex and birth weight, in full-term and preterm infants, and that deviations in predicted brain-age will correlate with health acuity.

## METHODS

### Study Populations

For the dHCP dataset (Edwards et al., 2022), quality control followed the exact steps outlined in https://github.com/EduNeuroLab/wmDevIuEu/blob/main/codeForMainFigures.ipynb (Grotheer et al., 2023). As a first quality assurance step, we removed all sessions that exceeded two standard deviations of the mean with regard to absolute motion and with regard to the amount of outlier slices replaced by FSL’s eddy tool during preprocessing. Five additional sessions were excluded during visual inspection due to obvious image artifacts or alignment errors. After tractography and bundle identification, we conducted an additional quality assurance by excluding all sessions where one or more bundles contained 10 or less streamlines. The remaining data consisted of 402 sessions from 368 individuals. The deduplicated dataset (n = 368) includes one scan per participant, with preference given to scans at later chronological ages when multiple scans were available for the same individual. We did not have health information about the dHCP dataset.

For the Lucile Packard Children’s Hospital dataset, participants for the clinical sample were 162 children born at <32 weeks gestational age (GA) who underwent magnetic resonance imaging (MRI) at near-term equivalent age. Infant clinical characteristics were collected from their Electronic Medical Records (EMR): gestational age (days), sex, birthweight (grams), and postmenstrual age (PMA; days) at time of MRI. We also collected data on major inflammatory morbidities associated with white matter brain injury: sepsis, bronchopulmonary dysplasia (BPD), necrotizing enterocolitis (NEC), and intraventricular hemorrhage (IVH). BPD was defined as treatment with supplemental oxygen at 36 weeks postmenstrual age (Feldman et al., 2002). Sepsis was defined as a positive blood culture or >7 days of antibiotics. IVH was defined as the presence of a grade I-III hemorrhage using the Papile classification system (Papile et al., 1978). NEC was defined as a diagnosis of medical or surgical NEC.

As detailed in Dubner et al. (2024), infants underwent magnetic resonance imaging (MRI) at the Stanford Lucile Packard Children’s Hospital as part of standard of care from Jun 2016 to Jan 2022. During the study period, MRI scans were conducted prior to hospital discharge for all infants born very preterm (<32 weeks GA) in our Neonatal Intensive Care Unit (NICU). Per clinical procedures, infants were scanned once stable in an open crib, requiring minimal respiratory support (no more than lowflow supplemental oxygen), and had reached a minimum postmenstrual age of 34 weeks. Before scanning, infants were fed, swaddled, and fitted with adhesive foam noise dampeners over their ears to block sounds from the scanner. Scans occurred during natural sleep with no sedation. MRI scans were acquired using one of two 3.0T scanners (GE Discovery MR750 or GE Signa Premier) equipped with either an 8-channel HD head coil or a 32-channel HD head coil (General Electric Healthcare, Little Chalfont, UK).

Sequences included in the clinical protocol and analyzed in this study included a high-resolution T1- weighted anatomical scan and 60-direction diffusion MRI scan. High-resolution T1-weighted scans were collected at approximately 0.430 x 0.430 x 1.400 mm^3^ spatial resolution. During preprocessing, T1-weighted images were resampled to 1.0mm³ isotropic voxels and used as infant-specific anatomical reference for diffusion tractography analyses. Diffusion MRI included a high-angular resolution (60-direction) single-shell sequence with a b-value= 700s/mm^2^ collected with a multi-slice echoplanar imaging (EPI) protocol for rapid image acquisition (∼3 min each) with two to six volumes at b=0. Diffusion MRI data were collected at 2.0mm^3^ spatial resolution (2.0mm x 2.0mm x 2.0mm isotropic voxels).

Subjects were excluded if they had a documented grade 4 intraventricular hemorrhage in either hemisphere, and if their scan had one of the following: mean framewise displacement > 2mm OR T1 neighbor correlation < 0.9mm OR raw neighbor correlation < 0.9mm. Subjects were also excluded if they had more than 6 tracts missing from tract segmentation, and if their PMA was greater than 3 standard deviations from the mean of the remaining sample. The total number of excluded subjects was 44, and 162 subjects remained in the final dataset after quality control.

Demographic variables were balanced across sexes in the sample with the exception of birthweight, and those that failed preprocessing also did not differ significantly from those who had neuroimaging data that completed preprocessing.

### Diffusion Data Preprocessing

Preprocessing for the LPCH dataset was performed using *QSIPrep* 1.0.0rc2.dev0+g789be41.d20241119 (Cieslak et al., 2021), which is based on *Nipype* 1.9.1 (Gorgolewski et al., 2011, Esteban et al., 2025, RRID:SCR_002502). The additional flags used were infant, output-resolution 2, unringing-method mrdegibbs, and ignore fieldmaps.

The anatomical reference image was reoriented into AC-PC alignment via a 6-DOF transform extracted from a full Affine registration to the MNIInfant+1 template. A full nonlinear registration to the template from AC-PC space was estimated via symmetric nonlinear registration (SyN) using antsRegistration. Brain extraction was performed on the T1w image using SynthStrip (Hoopes et al., 2022) and automated segmentation was performed using SynthSeg (Billot et al., 2023, @synthseg2) from FreeSurfer version 7.3.1.

Any images with a b-value less than 100 s/mm^2^ were treated as a *b*=0 image. MP-PCA denoising as implemented in MRtrix3’s dwidenoise (Veraart et al., 2016) was applied with an auto-voxel window. When phase data were available, this was done on complex-valued data. After MP-PCA, Gibbs unringing was performed using MRtrix3’s mrdegibbs (Kellner et al., 2016). Following unringing, the mean intensity of the DWI series was adjusted so all the mean intensity of the b=0 images matched across each separate DWI scanning sequence. B1 field inhomogeneity was corrected using dwibiascorrect from MRtrix3 with the N4 algorithm (Tustison et al., 2010) after corrected images were resampled.

FSL’s eddy was used for head motion correction and Eddy current correction (Andersson & Sotiropoulos, 2016). Eddy was configured with a *q*-space smoothing factor of 10, a total of 5 iterations, and 1000 voxels used to estimate hyperparameters. A linear first level model and a linear second level model were used to characterize Eddy current-related spatial distortion. *q*-space coordinates were forcefully assigned to shells. Field offset was attempted to be separated from subject movement. Shells were aligned post-eddy. Eddy’s outlier replacement was run (Andersson et al., 2016). Data were grouped by slice, only including values from slices determined to contain at least 250 intracerebral voxels. Groups deviating by more than 4 standard deviations from the prediction had their data replaced with imputed values. Final interpolation was performed using the jac method.

Several confounding time-series were calculated based on the preprocessed DWI: framewise displacement (FD) using the implementation in *Nipype* (following the definitions by Power et al., 2014). The head-motion estimates calculated in the correction step were also placed within the corresponding confounds file. Slicewise cross correlation was also calculated. The DWI time-series were resampled to ACPC, generating a *preprocessed DWI run in ACPC space* with 2mm isotropic voxels. For more details of the pipeline, see <u>the section corresponding to workflows in</u> *<u>QSIPrep</u>*<u>’s documentation</u>.

### Reconstruction

Reconstruction for the LPCH dataset was performed using *QSIRecon* 1.0.0rc3.dev2+gf95211a.d20241206 (Cieslak, Camacho, et al., 2025; Cieslak, Irfanoglu, et al., 2025), which is based on *Nipype* 1.9.1 (Gorgolewski et al., 2011, Esteban et al., 2025, RRID:SCR_002502). The additional flags used were infant, output-resolution 2, and slight modifications to the ss3t_noACT pipeline using a custom yaml file.

#### Anatomical data for DWI reconstruction

Brainmasks from antsBrainExtraction were used in all subsequent reconstruction steps.

#### MRtrix3 Reconstruction

Multi-tissue fiber response functions were estimated using the dhollander algorithm. FODs were estimated via constrained spherical deconvolution (CSD, Tournier et al., 2004, 2008) using an unsupervised multi-tissue method (Dhollander et al., 2016, 2019). A single-shell-optimized multi-tissue CSD was performed using MRtrix3Tissue (https://3Tissue.github.io), a fork of MRtrix3 (Tournier et al., 2019). FODs were intensity-normalized using mtnormalize (Raffelt et al., 2017).

Many internal operations of *QSIPrep* and *QSIRecon* use *Nilearn* 0.10.1 (Abraham et al., 2014, RRID:SCR_001362) and *Dipy* (Garyfallidis et al., 2014). For more details of the pipeline, see <u>the</u> <u>section corresponding to workflows in *QSIRecon*’s documentation</u>.

For each whole-brain tractogram, we adapted the following parameters (Grotheer et al., 2023): number of streamlines: 2 million, algorithm: iFOD2, step size: default (1mm for derivatives resampled to 2mm isotropic), minimum length: default (30mm), maximum length: default (250mm), FOD amplitude stopping criterion: default (0.1), maximum angle: default (45 degrees).

### Tractometry

We applied the pyAFQ v1.3.6 pipeline to perform advanced tractometry analysis (Kruper et al., 2025). Using pyAFQ, we fit constrained spherical deconvolution (CSD) and used it as the fiber orientation distribution function for probabilistic tractography implemented in DIPY (Garyfallidis et al., 2014; Tournier et al., 2008). We used symmetric normalization (SyN) (Avants et al., 2008) diffeomorphic non-linear registration to register subjects to the University of North Carolina neonatal template (Shi et al., 2011). We used non-linear registration because the linear registration applied during preprocessing does not capture subtle local anatomical differences in brain anatomy that need to be taken into account in defining the trajectory of major white matter tracts. Twenty-six different white matter tracts were defined in template space based on a combination of inclusion and exclusion regions of interest (ROI). Twenty-four are previously defined in babyAFQ (Grotheer et al., 2022) with the additional inclusion of the forceps major and forceps minor. Each tract was divided into 100 nodes, resulting in a feature space of 5200 features per subject utilizing both fractional anisotropy (FA) and mean diffusivity (MD). Diffusion metrics were set to NaNs (and therefore treated as missing) if the tract comprised of 10 or less streamlines after segmentation. This step preceded the subject-level exclusion criteria of 6 or above missing tracts.These tractometry features were then used in brain-age prediction models as described below.

### Brain-Age Models

We evaluated neonatal brain age models using white matter microstructural features in two independent datasets. The developing Human Connectome Project (dHCP) dataset release 2 comprises healthy preterm and full-term infants (n = 368), and includes a small sub-sample (n = 44) of preterm infants born before 32 weeks gestational age (Edwards et al., 2022). For dHCP infants with more than one scan, preference was given to scans at the later chronological age. This large, high-quality, publicly available dataset serves as a useful benchmark to evaluate our modeling approach. To evaluate generalizability to clinical populations, we applied the same approach to routine clinical MRI scans from high-risk preterm infants in Lucile Packard Children’s Hospital’s neonatal intensive care unit (LPCH, n = 162).

We predicted infants’ postmenstrual age at scan (PMA) using principal components regression with least absolute shrinkage and selection operator (Lasso; Tibshirani, 1996). Models employed nested cross-validation with 5-fold splits repeated 40 times (200 total folds). Although hyperparameters were consistent across datasets, models were trained independently on each dataset.

In some of the models, we controlled for sex and birthweight as covariates using the CovariateRegressor class and used the residuals to predict PMA, given their associations with brain structure and degree of prematurity, as well as the aforementioned correlation with PMA.

This CovariateRegressor class was adapted from the ConfoundRegressor class from Snoek et al. (2019) and their open-source implementation (https://github.com/lukassnoek/MVCA/blob/master/analyses/confounds.py), and has since been implemented in the latest version of AFQ-Insight. This approach removes the linear effects of specified confounds from brain imaging features following established best practices for confound control in decoding analyses of neuroimaging data. The ConfoundRegressor operates by fitting separate linear regression models for each brain feature, where confound variables (or covariates) serve as predictors and individual features as dependent variables.

For each feature f, the model takes the form:

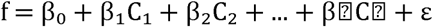

where CL through CL represent the covariates, βL is the intercept term, and ε represents the residual after covariate removal. The residuals from these regressions constitute the covariate-corrected features used in subsequent analyses.

Thereafter, for all predicted PMA values, we corrected them following equations 3 and 5 outlined in de Lange & Cole (2020) by first fitting the relationship between PMA predicted based on covariate-corrected white matter features in each participant and actual PMA:

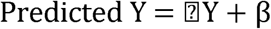

where Predicted Y is the predicted PMA from the residualized features that have been corrected for birthweight and sex, and Y is the actual PMA. The derived values of L and β are then used to correct PMA with

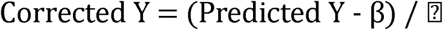

Finally, the corrected brain-age gap (cBAG) was calculated as the corrected predicted PMA (Corrected Y) minus actual PMA (Y), where negative gaps suggest delayed maturation and positive gaps indicate accelerated maturation.

### AFQ-Insight

Multidimensional analysis of informative features from tractometry was performed with AFQ-Insight 0.7.1 (Richie-Halford et al., 2021).

The PMA at scan was used as the target variable. A lasso principal components regression (using n-1 components) with elastic net regularization was used to predict targets, with splits used to ensure independence of training and test sets. We also included foldwise covariate regression to minimize bias as detailed in the Brain-Age Models section above. To evaluate model fit, we used a nested cross-validation procedure with 20% of the dataset held out for each batch, and predicted the age of held out subjects with fixed parameters based on the linear coefficients from the training set.

Our implementation extends the original MultiVoxel Confound Analysis (MVCA) framework with enhancements for cross-validation compatibility, missing data handling, preprocessing integration, numerical stability, and flexible data handling.

### Health Acuity Sum

Having established brain-age models for both datasets, we next examined clinical correlates in the LPCH sample. For the LPCH dataset, we examined four common conditions of prematurity: sepsis, necrotizing enterocolitis (both binarized as absent/present), bronchopulmonary dysplasia (grades 0-1 vs. 2+), and intraventricular hemorrhage (absent/grade 1 vs. grades 2-3; grade 4 in either hemisphere excluded). These were summed into an ordinal health acuity score, with a higher sum reflecting the presence of more conditions and thus a sicker infant.

### Bayesian Regression Models

We examined the relationship between brain-age gap and health outcomes using a series of ordinal Bayesian regression models with increasing complexity, where each parameter is estimated as a probability distribution rather than a point estimate:

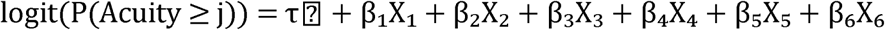

where Acuity represents the ordinal health acuity sums (levels 0, 1, 2, 3, 4), j indexes the acuity thresholds (j = 1, 2, 3), τ_l1_ are the threshold parameters in logits, X represents the predictor variables and β represents the mean of the posterior distribution for the various predictor variables. Positive coefficients increase the probability of being in higher acuity categories, and coefficients for the thresholds represent the log-odds of being at or above each acuity level when all predictors are equal to zero.

We used a cumulative model as any combination of health conditions could give rise to a higher health acuity sum (e.g. an infant with bronchopulmonary dysplasia will be assigned a sum score of 1, while an infant with sepsis, necrotizing enterocolitis, and intraventricular hemorrhage but not bronchopulmonary dysplasia will be assigned a sum score of 3). There were no infants with a sum score of 4 in this study.

Models were compared using Expected Log Pointwise Predictive Density (ELPD), where less negative scores indicate better model performance in predicting unseen test data during leave-one-out (LOO) cross validation. These models were fitted using Bambi release 0.15.0 (Capretto et al., 2022).

## RESULTS

### Study populations

While both datasets followed the same processing pipeline (Figure 1), they differ substantially in their clinical characteristics and study populations (Figure 2). The dHCP sample comprised primarily of full-term infants with a relatively small subsample of preterm infants. The LPCH sample, on the other hand, comprised high-risk premature infants who required extended medical care. The dHCP and LPCH datasets showed no significant difference in sex distribution, but the datasets were significantly different across all age-related measures (Table 1), including post-menstrual age at scan that was defined as the sum of gestational age at birth and chronological age at scan.

**Figure 1.**
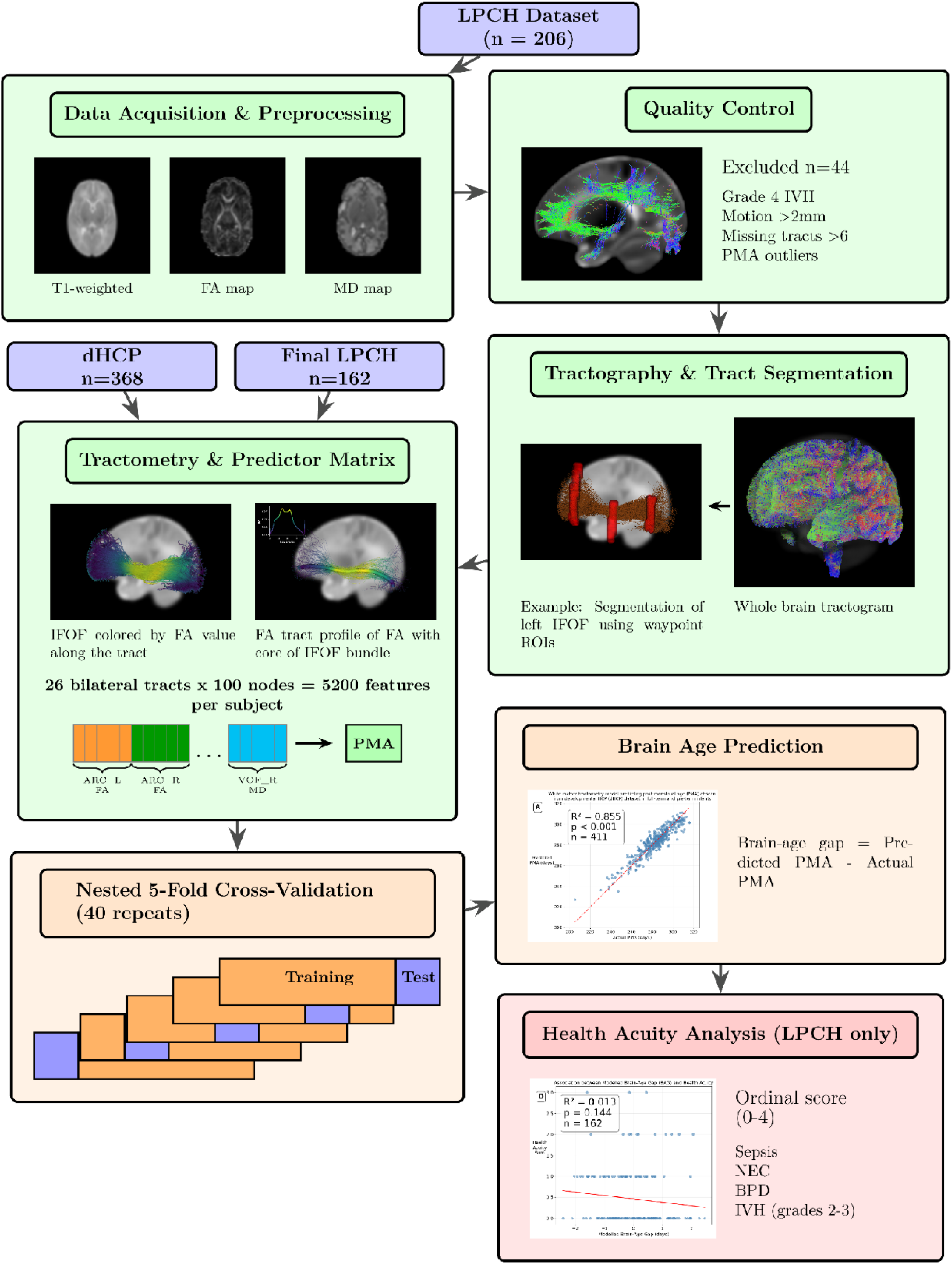
Study Design and Analysis Pipeline. The flowchart illustrates the complete data processing and analysis workflow for both datasets. The LPCH dataset (n=206) underwent data acquisition and preprocessing, followed by quality control (excluding n=44 subjects), tractography and tract segmentation (26 bilateral tracts), and tractometry (5,200 features per subject: 26 bilateral tracts × 100 nodes for both FA and MD). The dHCP dataset (n=368) was processed similarly, yielding final samples of LPCH n=162 and dHCP n=368. Each dataset was used independently to train separate brain-age prediction models using nested 5-fold cross-validation with 40 repeats (200 total folds per dataset). Postmenstrual age (PMA) served as the training target, with brain-age gap calculated as predicted minus actual PMA. For the LPCH sample only, brain-age gap was subsequently analyzed in relation to health acuity scores derived from four clinical complications (sepsis, necrotizing enterocolitis, bronchopulmonary dysplasia, and intraventricular hemorrhage grades 2-3), resulting in a health acuity ordinal score of 0-4.

**Figure 2.**
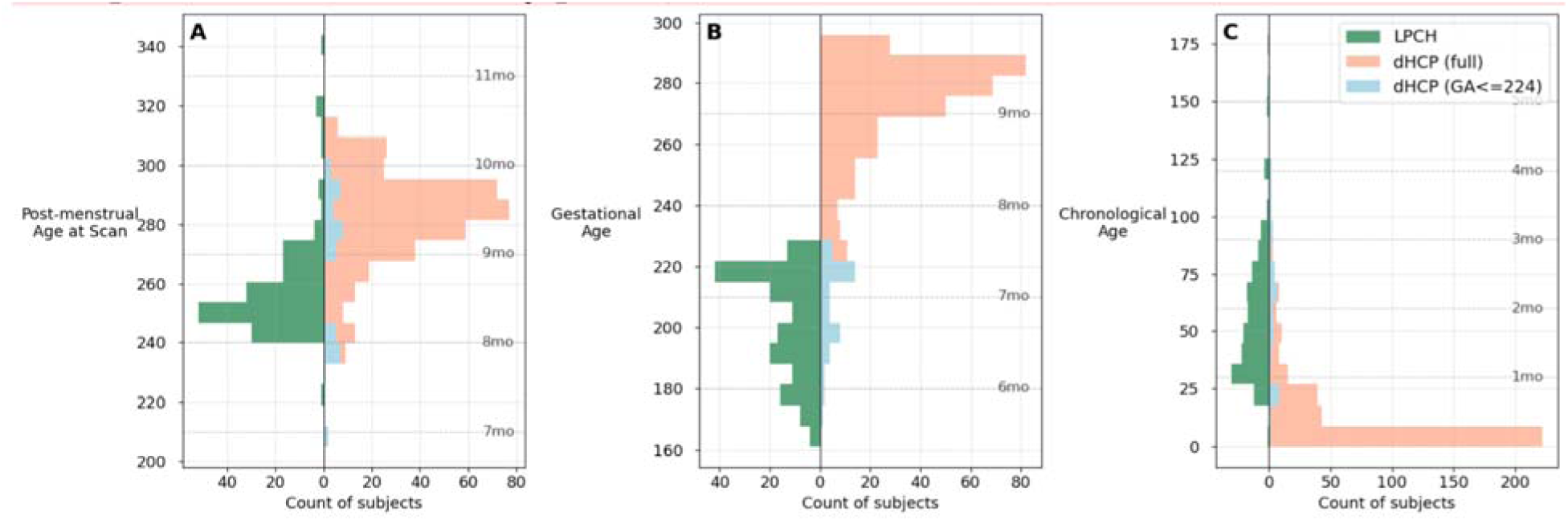
Age distribution comparison across both dHCP and LPCH datasets. Population pyramids showing age distributions for (A) post-menstrual age at scan, (B) gestational age at birth, and (C) chronological age at scan across both datasets. LPCH subjects (n = 162; green bars, left side) are compared with dHCP deduplicated dataset (n = 368; coral bars, right side) and dHCP subjects with gestational age ≤224 days (n = 44; light blue bars, right side). Dotted horizontal lines indicate monthly intervals. Ages are expressed in days.

**Table 1.**
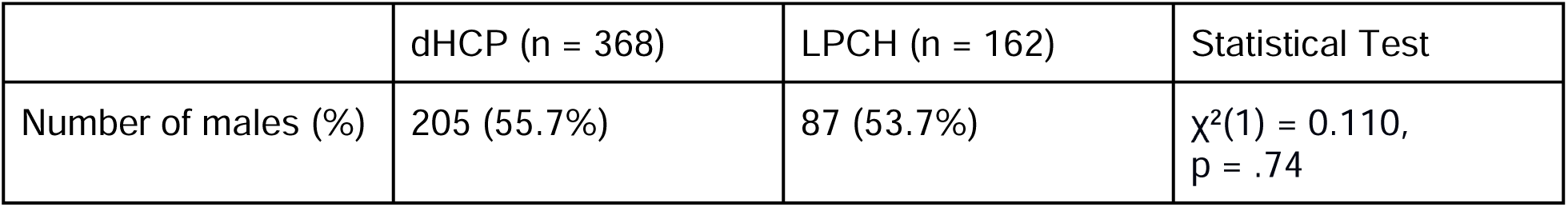
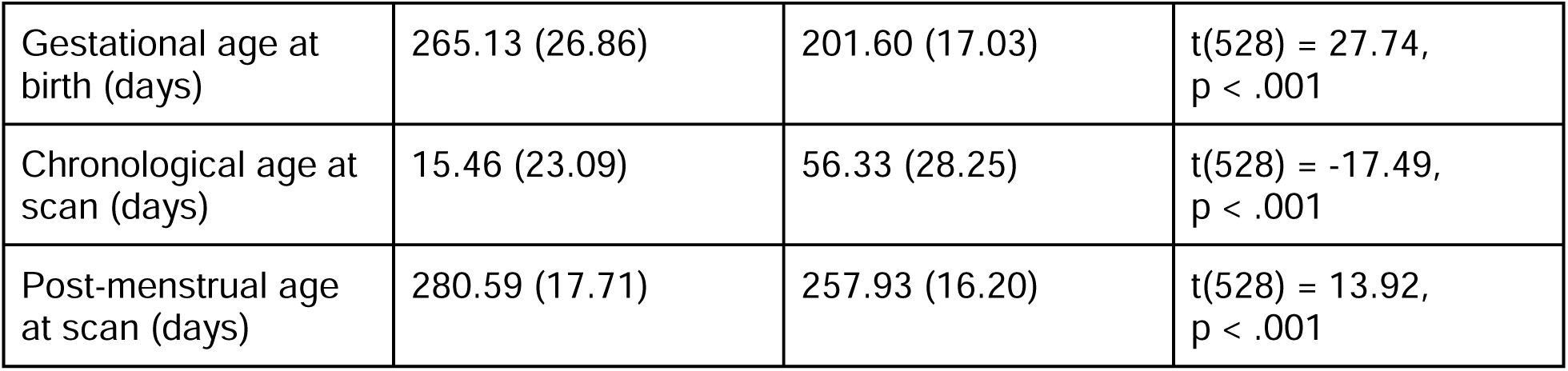
Demographic characteristics of study participants from dHCP and LPCH datasets. Data are presented as mean (standard deviation) for continuous variables and number (percentage) for categorical variables. Statistical comparisons between datasets were performed using chi-square tests for categorical variables and independent t-tests for continuous variables. dHCP = developing Human Connectome Project; LPCH = Lucile Packard Children’s Hospital; GA = gestational age; CA = chronological age; PMA = post-menstrual age. The datasets showed no significant difference in sex distribution (χ²(1) = 0.110, p = .74) but significant differences in all age-related measures: GA at birth (t(528) = 27.74, p < .001), CA at scan (t(528) = -17.49, p < .001), and PMA at scan (t(528) = 13.92, p < .001).

In both datasets, chronological age (CA) at scan was negatively correlated with gestational age (GA) and birthweight, with r = -0.76 and r = -0.65 respectively in the dHCP dataset, and r = - 0.86 and r = -0.68 respectively in the LPCH dataset. These negative correlations remained even after subsampling only very preterm infants in the dHCP dataset (r = -0.55 and r = -0.45, respectively). This reflects the clinical practice of delaying scans until infants achieve sufficient medical stability: the most preterm infants require prolonged hospital stays due to higher health acuity and are therefore scanned at older chronological ages prior to discharge.

### Brain age models in the dHCP dataset

We constructed brain-age models in the dHCP dataset to assess how well these features capture normative brain maturation across full-term and preterm infants. Table 2 demonstrates the performance of the brain age model of white matter features predicting post-menstrual age (PMA) at scan. There was minimal loss of predictive power between train and test splits with high coefficients of determination (R²) and mean absolute error less than 4 days.

**Table 2.**
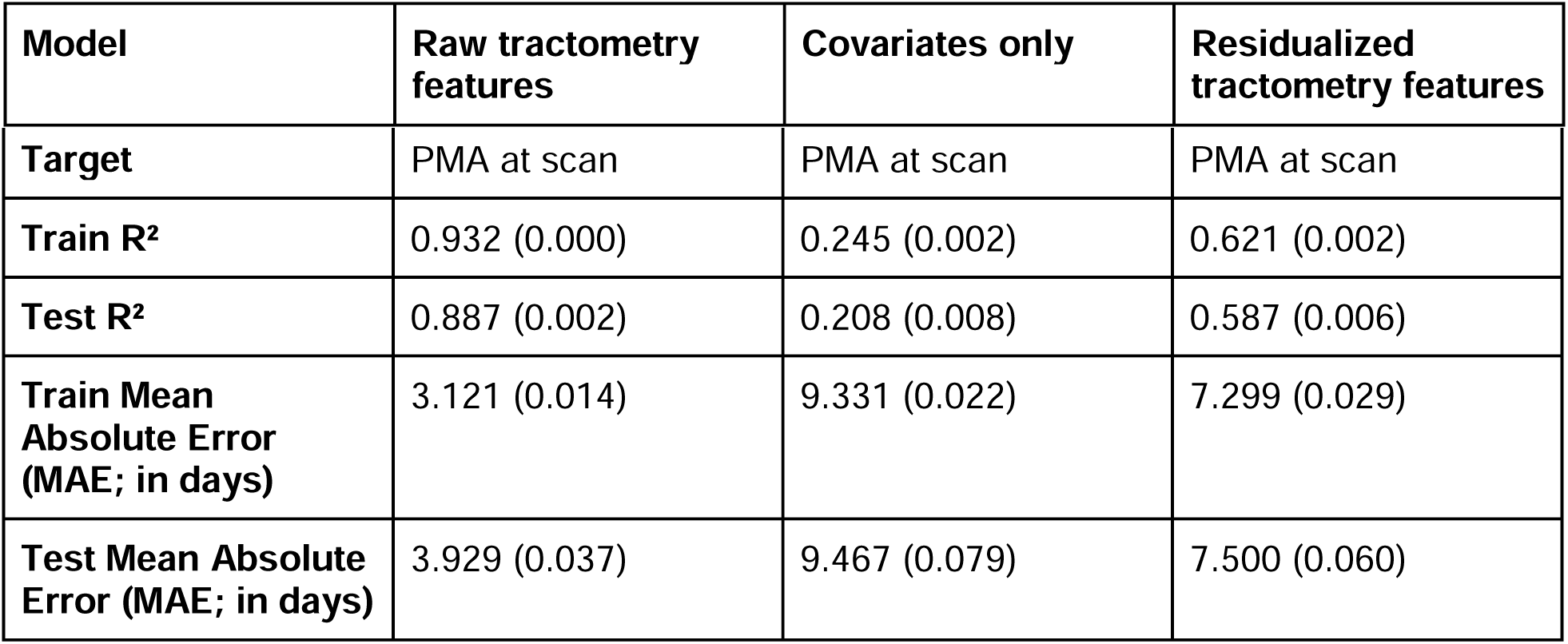
Model performance for predicting postmenstrual age at scan using white matter features in the dHCP dataset. Performance metrics are shown for three modeling approaches: raw tractometry features (with covariates included), covariates only (birthweight, sex), and residualized tractometry features (covariates regressed out). The deduplicated dataset (n = 368) includes one scan per participant, with preference given to scans at later chronological ages when multiple scans were available for the same individual. Values represent mean (standard error) across cross-validation folds. R² = coefficient of determination; MAE = mean absolute error in days; PMA = postmenstrual age.

The model constructed using raw white matter features achieved exceptional performance (Test R² = 0.887, MAE = 3.93 days), substantially outperforming the covariates-only model which utilized only birthweight and sex (Test R² = 0.208, MAE = 9.47 days). Notably, residualized white matter features retained considerable predictive power even after regressing out birthweight and sex covariates (Test R² = 0.587, MAE = 7.50 days), indicating that white matter contains information about maturation beyond these demographic factors. Covariates alone did not explain much variance in PMA at scan, but did augment the ability of white matter features to do so.

We then evaluated prediction of PMA at scan in the restricted dHCP sample consisting only of preterm infants born before 32 weeks gestation (very preterm) (Table 3). White matter features also demonstrated high accuracy (Test R² = 0.877, MAE = 5.40 days), becoming less accurate by 1.4 days compared to the full dHCP sample (MAE = 3.93 days), but still substantially outperforming the covariates-only model which showed poor generalization (Test R² = -0.317, MAE = 23.16 days), as well as the model using white matter features that controlled for covariates (Test R² = 0.765, MAE = 8.00 days). Together, these analyses indicate that tractometry metrics can form the basis of an accurate brain age model, surpassing the state of the art of 12.74 days of mean absolute error for preterm infants and 6.3 days for term infants using 3D convolutional neural networks (Fang et al., 2024). We next test how well this approach generalizes to high risk infants scanned in a clinical setting.

**Table 3.** Model performance for predicting postmenstrual age at scan using white matter features in very preterm infants from the dHCP dataset. Performance metrics are shown for three modeling approaches: raw tractometry features (with covariates included), covariates only (birthweight, sex), and residualized tractometry features (covariates regressed out). The very preterm subset (n = 44) includes only participants born before 32 weeks gestational age. Values represent mean (standard error) across cross-validation folds. R² = coefficient of determination; MAE = mean absolute error in days; PMA = postmenstrual age.

| Model | Raw tractometry features | Covariates only | Residualized tractometry features |
| --- | --- | --- | --- |
| Target | PMA at scan | PMA at scan | PMA at scan |
| Train R<sup>2</sup> | 0.956 (0.002) | 0.037 (0.003) | 0.857 (0.003) |
| Test R<sup>2</sup> | 0.877 (0.007) | -0.317 (0.055) | 0.765 (0.012) |
| Train Mean Absolute Error (MAE; in days) | 3.31 (0.091) | 21.13 (0.183) | 7.06 (0.108) |
| Test Mean Absolute Error (MAE; in days) | 5.40 (0.148) | 23.16 (0.314) | 8.00 (0.230) |

### Brain age models in the preterm clinical LPCH dataset

We evaluated prediction of PMA at scan in the LPCH dataset (Table 4). White matter features demonstrated moderate predictive performance for post-menstrual age at scan (Test R² = 0.391, MAE = 6.56 days), becoming less accurate by 1.2 days compared to the very preterm dHCP sample (Table 3; MAE = 5.49 days), but still outperforming the covariates-only model which included birthweight and sex (Test R² = 0.049, MAE = 7.97 days), as well as the model using white matter features that controlled for covariates (Test R² = 0.211, MAE = 7.64 days). Notably, in all cases, model performance was less accurate than in the dHCP dataset, likely reflecting differences in data quality, a restricted range of PMA, as well as the fact that all these infants had health complications.

**Table 4.**
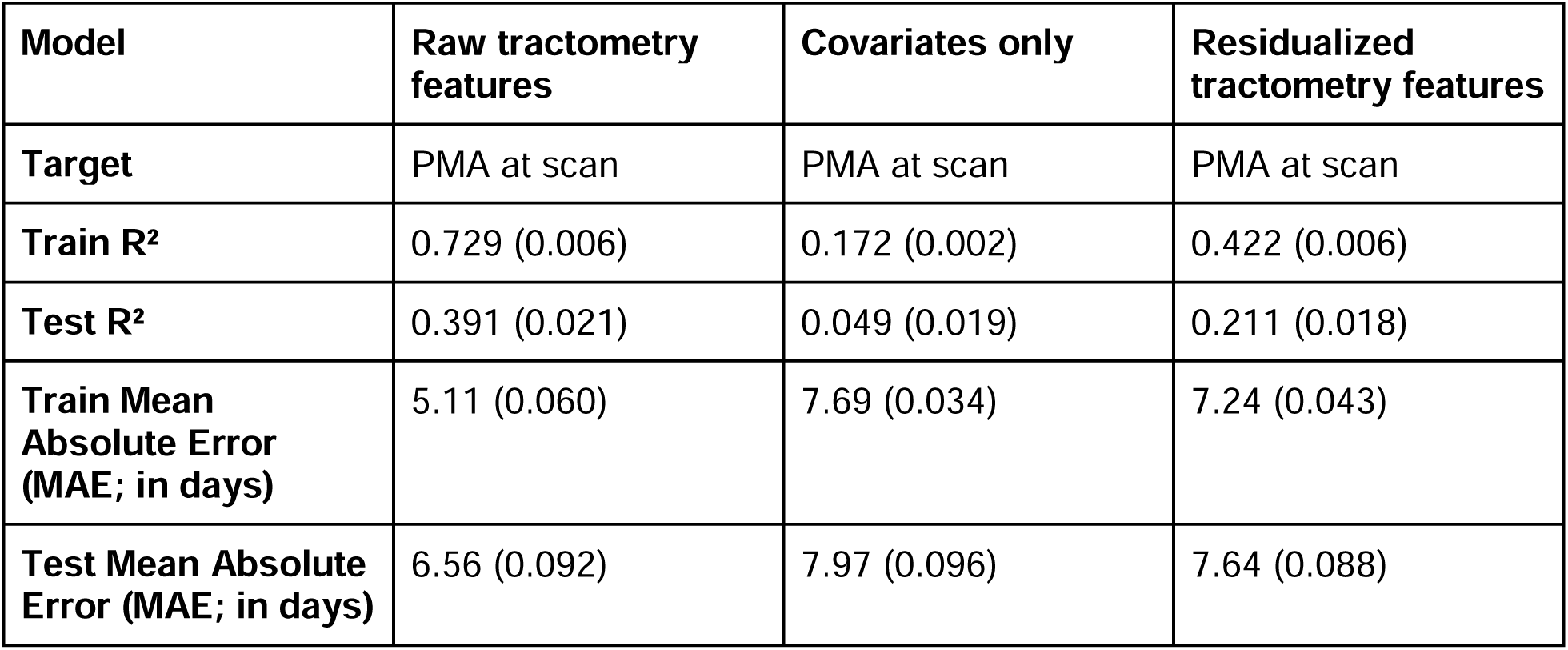
Model performance for predicting postmenstrual age at scan using white matter features in the preterm clinical dataset. Performance metrics are shown for three modeling approaches: raw tractometry features (with covariates included), covariates only (birthweight, sex), and residualized tractometry features (covariates regressed out). The LPCH dataset (n = 162) includes preterm infants scanned at a single clinical site. Values represent mean (standard error) across cross-validation folds. R² = coefficient of determination; MAE = mean absolute error in days; PMA = postmenstrual age.

| Model | Raw tractometry features | Covariates only | Residualized tractometry features |
| --- | --- | --- | --- |
| Target | PMA at scan | PMA at scan | PMA at scan |
| Train $R^2$ | 0.729 (0.006) | 0.172 (0.002) | 0.422 (0.006) |
| Test $R^2$ | 0.391 (0.021) | 0.049 (0.019) | 0.211 (0.018) |
| Train Mean Absolute Error (MAE; in days) | 5.11 (0.060) | 7.69 (0.034) | 7.24 (0.043) |
| Test Mean Absolute Error (MAE; in days) | 6.56 (0.092) | 7.97 (0.096) | 7.64 (0.088) |

### Association of brain-age gap with health in the preterm clinical dataset

A series of Bayesian ordinal regression models revealed that white matter-based brain-age gaps did not provide additional predictive information about health acuity in preterm infants beyond conventional clinical variables (n=161). Across all model specifications (see Supplementary Material), public insurance status showed the strongest association with health outcomes, while infants with public insurance had consistently lower illness severity.

Chronological age at scan was also positively associated with health acuity. This likely reflects clinical practice patterns in which scanning is delayed until infants achieve sufficient medical stability, such that the most preterm or acutely ill infants are scanned at older chronological ages prior to discharge. The corrected brain-age gap showed no credible associations with health outcomes in linear models, interaction models, or non-linear specifications. Model comparison using leave-one-out cross-validation confirmed that decomposing post-menstrual age into gestational age at birth and chronological age at scan provided the best predictive performance, while adding brain-age information did not improve model accuracy.

## DISCUSSION

Our study provides a comprehensive examination of the relationship between white matter microstructure, brain maturation, and health outcomes in preterm infants, utilizing advanced machine learning techniques and Bayesian modeling across two independent datasets: the developing Human Connectome Project (dHCP) and a clinical sample from Lucile Packard Children’s Hospital (LPCH). This investigation addresses three specific aims: constructing a neonatal brain-age model using tractometry-derived features, applying this model to routine clinical MRI data, and evaluating the model’s ability to provide health-related information beyond conventional measures.

Addressing our first aim, we successfully constructed an extremely accurate neonatal brain-age model using high-dimensional white matter features computed via tractometry. This model demonstrated strong predictive power for postmenstrual age (PMA) at scan in the dHCP dataset, which included both full-term and preterm infants, achieving accuracy equivalent to or better than state-of-the-art approaches using 3D convolutional neural networks (Fang et al., 2024). The model’s excellent performance supports our hypothesis that tractometry-derived features enable accurate brain-age prediction in this population that spans both full-term and preterm infants, reflecting rapid brain growth early in life. The success of white matter-based prediction likely reflects the particularly rapid and stereotyped microstructural changes occurring in white matter during the perinatal period, including progressive myelination and axonal development, processes that are associated with changes in diffusion properties accessible to dMRI measurements. Notably, tractometry effectively reduces the dimensionality of diffusion MRI data while preserving features relevant to individual differences to the extent that these differences are reflected in brain white matter tissue properties (Kruper et al., 2024), making it a computationally efficient yet powerful approach for brain-age modeling. Furthermore, we found that the model maintained its predictive ability, albeit with reduced performance, when applied to a sample of very preterm infants from the dHCP dataset. This suggests that our brain-age model is indeed applicable to preterm populations, though with some limitations that may reflect the unique developmental trajectories of very preterm infants.

The varying performance across datasets (dHCP, very preterm dHCP, and LPCH) underscores the critical importance of population-specific considerations in developmental neuroimaging.

This highlights the necessity of carefully controlling for demographic and clinical variables when interpreting brain imaging data in preterm populations. It emphasizes the need for cautious interpretation of age-related findings and underscores the pivotal role of covariate regression in isolating genuine age-related effects in white matter microstructure, especially when chronological age at scan and gestational age at birth are negatively correlated. These results collectively stress the importance of tailored approaches when studying diverse neonatal populations and the potential pitfalls of generalizing findings across different datasets without proper consideration of population-specific factors.

For the very preterm subsample of the dHCP dataset, the confound-only model performed poorly (negative cross-validated R^2^) but the unresidualized white matter models still outperformed the model using residualized features. This apparent discrepancy is again explained by the role of shared variance: although birthweight and sex had little standalone predictive value in the subsampled dHCP dataset, their covariance with white matter features contributed a biologically meaningful signal. When brain features were residualized for these confounds, the shared signal was removed, reducing model performance. In the restricted dHCP dataset, confounds contribute to prediction not in isolation but through their overlap with variation in white matter features.

Addressing our second aim, we successfully constructed a similar brain-age model using routine clinical MRI data from the LPCH dataset. While the performance was reduced compared to the dHCP dataset, the model still demonstrated moderate predictive power. This achievement highlights the potential for translating advanced neuroimaging techniques to clinical settings.

However, several factors likely contribute to the reduced performance in the clinical dataset. First, the LPCH data reflect routine clinical acquisitions with greater variability in imaging parameters compared to the standardized research protocols of dHCP. Second, the datasets differ in population characteristics, with potential differences in demographic composition and health acuity distributions. For example, dHCP includes both full-term and preterm infants (23-44 weeks’ GA), while LPCH comprises only preterm infants with more extremely preterm infants.

Regarding our third aim, we evaluated whether our preterm white matter-based brain-age model provided additional information about infant health beyond conventional clinical and demographic measures. Contrary to our hypothesis, we found no significant association between brain-age gap and health complications in the LPCH dataset. We observed only a slight negative trend suggesting that infants with more developmentally advanced brains relative to their chronological age may experience fewer health complications though, based on Bayesian regression models, this effect was not statistically credible. Thus, white matter microstructural features at term-equivalent age may not retain detectable variance of the cumulative burden of common prematurity-related complications, at least in the clinical data acquired here.

It is important to note that the clinical factors examined in this study (IVH, BPD, NEC, and sepsis) represent only a subset of influences on preterm brain development. Other potentially modifiable factors that may also drive white matter maturation, such as developmental care practices, nutrition, and cumulative pain and stress exposure, represent important avenues for future investigation. The current lack of strong associations with major structural morbidities (while modifiable care factors remain unexamined) may actually be encouraging - it would suggest that intervention opportunities exist beyond managing acute complications.

The brain-age gap measure, which uses information from all white matter tracts, might give us a more complete picture of brain development than looking at specific regions alone. However, even with this increased precision from a whole-brain approach, the brain-age gap didn’t show a significant link to health acuity. This suggests that the relationship between brain structure and health in preterm infants might be more complicated than what we can measure with current diffusion MRI techniques and linear models. It is important to consider what this global maturation metric actually captures. While preterm birth is associated with well-documented brain injury patterns (such as pre-oligodendrocyte vulnerability to hypoxia-ischemia documented in animal models), our tractometry-based brain-age model may primarily reflect the ongoing organizational and maturational processes of surviving tissue rather than specific injury or inflammatory markers. In this framework, white matter development may continue along a normative trajectory despite acute insults, with brain age capturing the health and developmental tempo of intact tissue.

In conclusion, our study provides valuable insights into the relationship between white matter development, brain maturation, and health outcomes in preterm infants. We successfully constructed a neonatal brain-age model using tractometry-derived features, demonstrated its applicability to routine clinical MRI data, and explored its potential to provide health-related information beyond conventional measures. The lack of association between brain-age gaps and health acuity suggests that global white matter maturation may be insufficiently sensitive to the cumulative burden of NICU-related morbidities, underscoring the need for multimodal or longitudinal biomarkers. Future work incorporating fetal MRI data (such as the HEALthy Brain and Child Development (HBCD) Study; Nelson et al., 2024) could provide a more refined developmental reference. By capturing brain maturation in utero without the confounding influences of extrauterine adaptation, such datasets would provide a purer comparison for assessing the impact of preterm birth on brain development trajectories.

Our study has laid important groundwork for future investigations in this field. The ability to accurately predict brain maturation from MRI data holds promise for clinical applications, potentially aiding in the identification of infants at risk for developmental delays or other complications. However, the lack of strong associations between brain-age gaps and health outcomes suggests that the clinical utility of these measures may be limited in their current form. As we continue to refine our methods and expand our understanding of preterm brain development, we move closer to the goal of improving outcomes for this vulnerable population through early identification of risk and targeted interventions.

## DATA AND CODE AVAILABILITY

Data for this paper is available at https://doi.org/10.6084/m9.figshare.c.7847159. Code for this paper is available at https://github.com/chiuhoward/neonatalbrainage.

## AUTHOR CONTRIBUTIONS

Howard Chiu

Conceptualization, Data Curation, Formal Analysis, Investigation, Methodology, Software, Visualization, Writing - original draft, Writing - review and editing.

Adam C. Richie Halford

Formal Analysis, Methodology, Software, Supervision, Writing - review and editing.

Molly F. Lazarus

Data Curation, Investigation, Project Administration, Writing - review and editing.

Ariel Rokem

Conceptualization, Methodology, Resources, Software, Validation, Writing - review and editing.

Rocio Velasco Poblaciones

Data Curation, Writing - review and editing.

Virginia A. Marchman

Data Curation, Formal Analysis, Writing - review and editing.

Katherine E. Travis

Conceptualization, Funding Acquisition, Investigation, Project Administration, Validation, Writing - review and editing.

Melissa L. Scala

Supervision, Writing - review and editing.

Heidi M. Feldman

Conceptualization, Funding Acquisition, Resources, Supervision, Writing - review and editing.

Jason D. Yeatman

Conceptualization, Funding Acquisition, Methodology, Project Administration, Resources, Supervision, Writing - review and editing.

## FUNDING

Howard Chiu is graciously funded by the Stanford Graduate Fellowship. This work was supported by the following grants from the Eunice Kennedy Shriver National Institute of Child Health and Human Development: 5R00-HD8474904 (K.E. Travis, PI), 2R01-HD069150 (H.M. Feldman, P), R01HD095861, and R01HD116845 (J.D. Yeatman, PI). Software development was also supported through R01EB027585 (PI: Eleftherios Garyfallidis, Indiana University, Rokem, sub-contract PI).

## DECLARATION OF COMPETING INTERESTS

The authors declare no potential conflicts of interest with respect to the research, authorship, and publication of this article.

## ACKNOWLEDGEMENTS

We would also like to thank all the families and healthcare professionals that made this research possible.

## DECLARATION OF GENERATIVE AI AND AI-ASSISTED TECHNOLOGIES IN THE MANUSCRIPT PREPARATION PROCESS

During the preparation of this work the authors used Claude in order to assist copy editing of this manuscript, and Gemini Deep Research in order to identify additional journal articles that may be relevant. After using these tools/services, the authors reviewed and edited the content as needed and take full responsibility for the content of the published article.

## SUPPLEMENTARY MATERIAL

Socioeconomic status (SES) is a powerful predictor of neurodevelopmental outcomes in preterm infants, with lower SES predicting alterations in brain development including growth of cerebral cortex and subcortical structures. The association between brain injury and poorer cognition is attenuated in children born to mothers of higher education level, suggesting that higher SES provides protective factors that can mitigate the impact of neonatal brain injury.

We also controlled for male sex in the linear models as male sex is a well-established risk factor for poor neurodevelopmental outcome after premature birth, with males showing higher mortality and poorer long-term neurologic outcomes compared to females, even controlling for gestational age and nutritional factors.

Unexpectedly, public insurance, a proxy for lower SES, was associated with lower health acuity scores in the LPCH dataset. This finding contrasts with prior literature linking lower SES to worse neonatal outcomes (Johnson et al., 2023; Thomson et al., 2021). The continued robust association between chronological age at scan and health acuity indicates that hospital length of stay remains the strongest predictor of illness severity in this population. One plausible explanation lies in the referral and case-mix characteristics of a tertiary care center. Privately insured infants may be disproportionately referred to LPCH for complex or rare conditions that inflate their health acuity scores, whereas publicly insured infants may represent a more routine preterm population. Additionally, differences in length of hospitalization, with medically complex, privately insured infants potentially remaining hospitalized longer, may have further elevated their health acuity scores. This pattern underscores the importance of considering institutional referral patterns, case-mix composition, and the actual operationalization of “health acuity” when interpreting SES-health associations in NICU populations.

---

**Table S1.** Baseline Bayesian ordinal regression model predicting health acuity sum using covariates only in the LPCH dataset. Health acuity sum represents a composite measure of illness severity (range 0-4; observed range 0-3) in preterm infants (n=161; one infant did not have birthweight recorded). Parameters show mean posterior estimates with standard deviations (SD). Highest Density Interval (HDI) represents the 94% credible interval; parameters with HDI ranges excluding zero indicate credible effects. R-hat values of 1.0 indicate excellent model convergence.

| Base model (covariates only): |  |  |  |  |  |
| --- | --- | --- | --- | --- | --- |
| health_acuity_sum ~ birthweight + male_sex + public_insurance |  |  |  |  |  |
| Parameter | Mean | SD | HDI 3% | HDI 97% | R-hat |
| birthweight | 0.089 | 0.160 | -0.196 | 0.401 | 1.0 |
| male_sex | -0.416 | 0.314 | -0.996 | 0.183 | 1.0 |
| Public insurance | -1.096 | 0.343 | -1.714 | -0.438 | 1.0 |

**Table S2.** Bayesian regression model predicting health acuity sum including post-menstrual age at scan in the LPCH dataset. Health acuity sum represents a composite measure of illness severity (range 0-4; observed range 0-3) in preterm infants (n=161; one infant did not have birthweight recorded). Parameters show mean posterior estimates with standard deviations (SD). Highest Density Interval (HDI) represents the 94% credible interval; parameters with HDI ranges excluding zero indicate credible effects. R-hat values of 1.0 indicate excellent model convergence.

| PMA model: |  |  |  |  |  |
| --- | --- | --- | --- | --- | --- |
| health_acuity_sum ~ post_menstrual_age_at_scan + birthweight + male_sex + public_insurance |  |  |  |  |  |
| Parameter | Mean | SD | HDI 3% | HDI 97% | R-hat |
| Post-menstrual age at scan | 0.881 | 0.185 | 0.525 | 1.216 | 1.0 |
| birthweight | 0.111 | 0.174 | -0.214 | 0.435 | 1.0 |
| male_sex | -0.477 | 0.337 | -1.127 | 0.138 | 1.0 |
| Public insurance | -1.037 | 0.353 | -1.672 | -0.351 | 1.0 |

**Table S3.** Bayesian ordinal regression model predicting health acuity sum with decomposed age variables in the LPCH dataset. Post-menstrual age at scan was decomposed into gestational age at birth and chronological age at scan. Health acuity sum represents a composite measure of illness severity (range 0-4; observed range 0-3) in preterm infants (n=161; one infant did not have birthweight recorded). Parameters show mean posterior estimates with standard deviations (SD). Highest Density Interval (HDI) represents the 94% credible interval; parameters with HDI ranges excluding zero indicate credible effects. R-hat values of 1.0 indicate excellent model convergence.

| CA and GA model: |  |  |  |  |  |
| --- | --- | --- | --- | --- | --- |
| health_acuity_sum ~ gestational_age_at_birth + chronological_age_at_scan + birthweight + male_sex + public_insurance |  |  |  |  |  |
| Parameter | Mean | SD | HDI 3% | HDI 97% | R-hat |
| Gestational age at birth | -0.466 | 0.332 | -1.104 | 0.126 | 1.0 |
| Chronological age at scan | 0.911 | 0.334 | 0.282 | 1.533 | 1.0 |
| birthweight | 0.067 | 0.186 | -0.278 | 0.416 | 1.0 |
| male_sex | -0.534 | 0.346 | -1.170 | 0.142 | 1.0 |
| Public insurance | -1.151 | 0.386 | -1.889 | -0.452 | 1.0 |

**Table S4.**
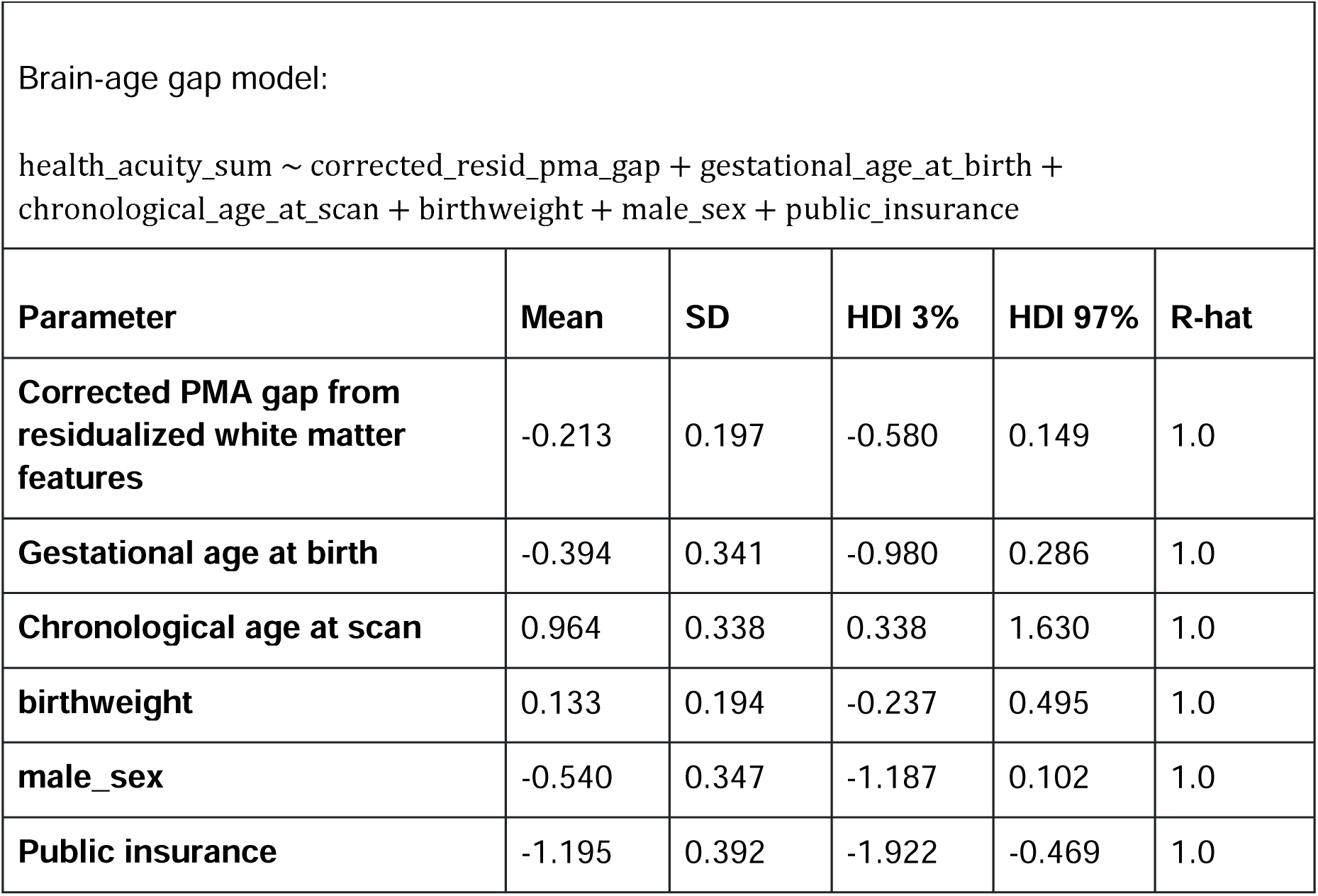
Bayesian ordinal regression model predicting health acuity sum including brain-age gap derived from white matter features in the LPCH dataset. The corrected brain-age gap represents the difference between corrected predicted PMA calculated using residualized white matter features that control for birthweight and sex, and actual PMA (in days). Health acuity sum represents a composite measure of illness severity (range 0-4; observed range 0-3) in preterm infants (n=161; one infant did not have birthweight recorded). Parameters show mean posterior estimates with standard deviations (SD). Highest Density Interval (HDI) represents the 94% credible interval; parameters with HDI ranges excluding zero indicate credible effects. R-hat values of 1.0 indicate excellent model convergence.

| Parameter | Mean | SD | HDI 3% | HDI 97% | R-hat |
| --- | --- | --- | --- | --- | --- |
| **Corrected PMA gap from residualized white matter features** | -0.213 | 0.197 | -0.580 | 0.149 | 1.0 |
| **Gestational age at birth** | -0.394 | 0.341 | -0.980 | 0.286 | 1.0 |
| **Chronological age at scan** | 0.964 | 0.338 | 0.338 | 1.630 | 1.0 |
| **birthweight** | 0.133 | 0.194 | -0.237 | 0.495 | 1.0 |
| **male_sex** | -0.540 | 0.347 | -1.187 | 0.102 | 1.0 |
| **Public insurance** | -1.195 | 0.392 | -1.922 | -0.469 | 1.0 |

**Table S5.** Bayesian ordinal regression model predicting health acuity sum including brain-age gap with interaction and quadratic terms in the LPCH dataset. This model included quadratic and interaction terms to test for non-linear relationships and moderation effects. The corrected brain-age gap represents the difference between corrected predicted PMA calculated using residualized white matter features that control for birthweight and sex, and actual PMA (in days). Health acuity sum represents a composite measure of illness severity (range 0-4; observed range 0-3) in preterm infants (n=161; one infant did not have birthweight recorded). Parameters show mean posterior estimates with standard deviations (SD). Highest Density Interval (HDI) represents the 94% credible interval; parameters with HDI ranges excluding zero indicate credible effects. R-hat values of 1.0 indicate excellent model convergence.

| Parameter | Mean | SD | HDI 3% | HDI 97% | R-hat |
| --- | --- | --- | --- | --- | --- |
| **(Corrected PMA gap)<sup>2</sup>** | 0.027 | 0.154 | -0.264 | 0.310 | 1.0 |
| **Corrected PMA gap from residualized white matter features** | -0.206 | 0.208 | -0.607 | 0.168 | 1.0 |
| **Gestational age at birth** | -0.538 | 0.373 | -1.246 | 0.154 | 1.0 |
| **Corrected PMA gap:GA** | -0.369 | 0.381 | -1.131 | 0.301 | 1.0 |
| **Chronological age at scan** | 0.854 | 0.361 | 0.207 | 1.552 | 1.0 |
| **Corrected PMA gap:CA** | -0.325 | 0.407 | -1.088 | 0.437 | 1.0 |
| **birthweight** | 0.128 | 0.195 | -0.237 | 0.492 | 1.0 |
| **male_sex** | -0.548 | 0.361 | -1.228 | 0.101 | 1.0 |
| **Public insurance** | -1.215 | 0.391 | -1.960 | -0.512 | 1.0 |

## Notes

https://github.com/chiuhoward/neonatalbrainage

https://doi.org/10.6084/m9.figshare.c.7847159

### Competing Interest Statement

The authors have declared no competing interest.

